# Measuring the information budget of cross-target antibody affinity generalization: a robust cross-target wall, and why structure reaches developability but not affinity

**DOI:** 10.64898/2026.11.17.767172

**Authors:** Luciano Martins, Priscila Pini Zenatti, João Meidanis

## Abstract

Structure-based machine learning is widely expected to advance antibody engineering, yet it is unclear how far affinity predictions transfer to new targets, and why. We introduce a noise-corrected, censoring-aware estimator of the cross-target transfer ceiling, *ρ*_max_ = 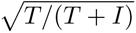, which separates a transferable antibody-intrinsic component (*T*) from a non-transferable target-specific interaction (*I*). On deterministic synthetic ground truth the estimator recovers the noise-free ceiling to *±*0.0003, whereas the naive sum-of-squares estimator is biased downward by measurement noise. Applying it from synthetic to real systems, the transferable budget is small, and on real neutralization panels it is not an artefact: the target-specific interaction survives noise and censoring correction. It is strongly regime-dependent, with a two-tier structure: predicting a *novel* antibody from sequence alone (the design setting) reaches a lower ceiling (*ρ ≈* 0.44 on real breadth data) than ranking a *characterized* one (*ρ ≈* 0.59), even with a strong protein language model. Synthetic oracles also overstate transferability: an oracle scored on real predicted structures carries about half the budget of a matched synthetic lattice (0.45 vs. 0.81), and real neutralization and escape panels sit well below the synthetic ceiling (0.32–0.68). Under leakage-free antigen-cluster splits, cross-target affinity ranking across every method family collapses to this budget, while within a sampled target a structure-based graph network ranks affinity above off-the-shelf FoldX and GearBind (within-complex Spearman 0.55), and structural surface descriptors predict a surface-driven developability property (hydrophobic aggregation) where sequence-dominated properties (humanness, charge-driven self-association) do not benefit. A controlled ablation attributes the residual within-target signal to the antibody alone: dropping the antigen from the structural model leaves it essentially unchanged (interface Δ*ρ* between *−*0.01 and +0.02 across four panels), so for this model the paratope-epitope interface adds no ranking signal over the antibody alone. Models climb toward the budget within a target—via a measure-then-rank operating mode that recovers top binders 6–8*×* more efficiently than random once a target has been sampled—but none crosses it.

## 1 Introduction

Monoclonal antibodies are among the fastest-growing classes of biopharmaceuticals [1, 2], and binding affinity, quantified by the dissociation constant *K_d_*, is the property that engineering campaigns most often seek to predict and optimize [3]. The maturation of protein structure prediction [4, 5] and of geometric deep learning [6, 7] has made structure-based affinity prediction appealing: a model that could rank candidate antibodies for a target from structure would reduce the experimental cost of discovery.

A broad spectrum of methods now targets antibody–antigen affinity. Physics-based force fields such as FoldX estimate the change in binding free energy (ΔΔ*G*) from a complex structure [8]; supervised geometric networks such as GearBind learn ΔΔ*G* from interface graphs with contrastive pretraining [7]; graph-signature predictors such as mCSM-AB2 score single-point substitutions [9]; cofolding models (AlphaFold 3, Boltz-2) predict the complex and, increasingly, a confidence or affinity signal [4, 5]; and sequence models (ESM-2 [10] and antibody-specific language models such as AntiBERTy [11]) predict affinity or its changes without explicit structure. Inverse-folding likelihoods and retrieval provide further structure-aware but training-free baselines. These methods are typically evaluated on mutational series within a small number of complexes, a regime that rewards memorizing target-specific signal.

The evaluation regime is where the difficulty begins. Volkov et al. showed for protein–ligand affinity that random splits, which leak targets between train and test, dramatically overstate generalization [12], and the same pathology affects antibody benchmarks [13]. Recent work documents the symptom: generalizable ΔΔ*G* prediction requires far more diverse data [14], and sequence models can match or beat structure on unseen antigens [13]. What these do not do is isolate the *cause* (whether cross-target failure reflects imperfect predicted structures or an intrinsic, informational limit), nor place the affinity result within the broader question of which antibody properties structure can predict at all. We close both gaps here.

We address this using a unified, decontaminated corpus of experimentally measured *K_d_* spanning 2,015 antigens, released as a companion data descriptor [15], together with established benchmarks [16, 17], synthetic ground truth [18], and real crossed antibody–antigen panels (HIV neutralization and SARS-CoV-2 escape). We map where structure helps and where it stops. (i) A quantitative boundary: structure helps surface-geometry developability (hydrophobic aggregation) and single-complex mutational ranking, but not cross-target affinity, not the antibody-antigen interface for this model (a controlled ablation shows it adds nothing to within-target ranking beyond the antibody alone), and not sequence-dominated properties (humanness, charge-driven self-association). (ii) The cross-target boundary is *informational*, not representational, and we *measure* it: a noise-corrected, censoring-aware estimator of the transfer ceiling *ρ*_max_ = 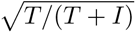, validated on deterministic synthetic ground truth (it recovers the noise-free ceiling to *±*0.0003 while the naive estimator is noise-biased), quantifies the non-transferable target-specific component across a ladder of synthetic and real systems; the budget is small, real on the neutralization and escape panels (it survives noise and censoring correction), two-tier (a *novel* antibody predicted from sequence transfers worse than a *characterized* one), and overstated by synthetic oracles. Supporting this map are (iii) a within-target structure advantage over sequence and physics baselines, (iv) a cross-target collapse reproduced across every method family under identical leakage-free antigen-cluster splits (plus physics, inverse-folding and MM-GBSA baselines), and (v) a practical consequence, measure-then-rank, for how these models should be deployed. All evaluations use antigen-grouped, leakage-free splits.

## 2 Results

### 2.1 Structure improves within-target affinity ranking

We first ask whether structure helps in the regime it is most likely to: ranking variants of a target for which measurements already exist. On SKEMPI 2.0 [16], a structure-based delta-siamese graph neural network (a geometric-vector-perceptron network operating on predicted complex graphs; see Methods) attains a within-complex Spearman correlation of **0.548**, ahead of GearBind with pretraining (0.465) [7], FoldX (0.436), and GearBind (0.404). Against a sequence-only model of matched capacity the structure edge is small and confined to single-antibody crystal panels: on a position-held-out split (which removes the near-duplicate leakage of a random within-mutant split) the gain is only +0.04 to +0.05 Spearman, and on genuinely diverse Boltz-predicted lineages it is null (mean +0.005, structure favored in 2*/*5; *p* = 0.95), consistent with the structure-quality dependence (Supplementary Note 5): structure improves within-target ranking only for single-antibody mutational series on accurate structures. The larger +0.093 figure reported in earlier drafts reflected a random within-mutant split and does not survive a leakage-free position split.

On the AbBiBench crystal set [19] the structure model correlates far more strongly than FoldX (*|ρ| ≈* 0.42 vs. 0.11 per-complex Spearman). This is a measure-then-rank comparison: the structure model is trained on target-specific measurements, whereas FoldX and GearBind are applied off-the-shelf (Figure 1).

**Figure 1:**
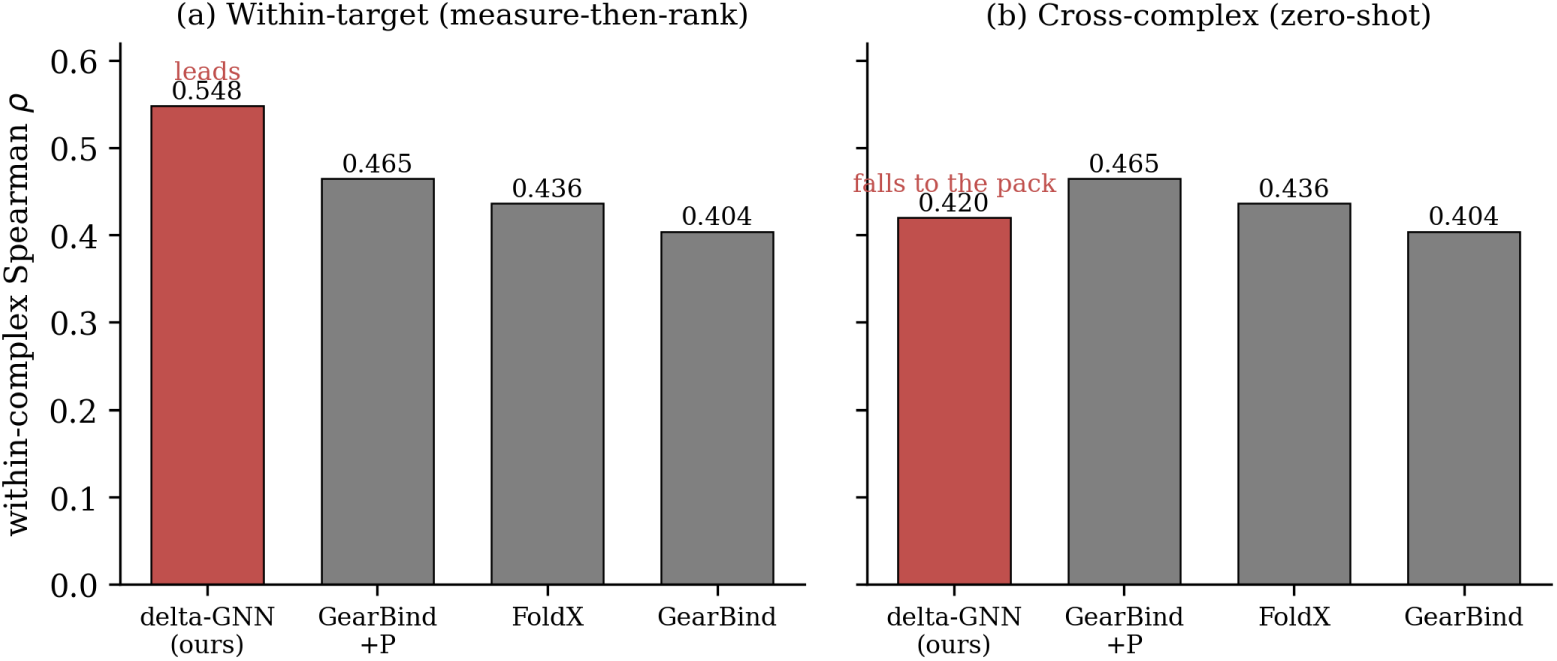
Structure ranks affinity within a target but not across targets (SKEMPI, 35 complexes, within-complex Spearman). (a) In the within-target (measure-then-rank) regime the structure-based delta-GNN leads (0.548) over off-the-shelf GearBind+pretrain (0.465), FoldX (0.436), and GearBind (0.404). (b) In the cross-complex (zero-shot) regime the same model falls to the pack (0.420), while the off-the-shelf baselines, being split-invariant, are unchanged. The delta-GNN receives target-specific training in (a) but not in (b); the baselines are off-the-shelf throughout.

The advantage depends on structure *quality* : FoldX, computed on predicted rather than crystallographic complexes, collapses to *ρ ≈* 0 (Supplementary Note 5). Structure is thus informative for within-target affinity, but only when the structure is good (Supplementary Note 5).

The advantage is also regime-specific. In a complementary cross-complex scenario, in which an entire complex is withheld, the structure model no longer leads: its within-complex ranking falls to 0.42, mid-pack among the off-the-shelf baselines (FoldX 0.44, GearBind 0.40, GearBind with pretraining 0.47), an early sign of the cross-target wall of Section 2.2. We further note that a widely used graph-signature predictor, mCSM-AB2 [9], cannot be benchmarked fairly in this setting: it scores only single-point substitutions, whereas the benchmarks are dominated by multi-point mutants, and its training set overlaps the antibody complexes used here, so its high apparent accuracy reflects memorization rather than generalization (Supplementary Note 2).

### 2.2 No method generalizes to unseen antigens: a cross-target wall

The first boundary is across targets. We evaluate generalization to *new* antigens under a strict, leakage-free protocol: antigen sequences are clustered with MMseqs2 (1,752 antigens *→* 549 clusters) and folds are grouped by cluster, so that near-duplicate antigens (for instance the SARS-CoV-2 spike and its receptor-binding domain, which would otherwise leak across folds and inflate apparent generalization) are always held out together. On GRABS we run a five-fold cluster leave-one-antigen-out split (4,020 variants) and, because a model can inflate a pooled *R*^2^ simply by predicting each antigen’s mean offset, we report the honest ranking metric: the *within-held-out-antigen* Spearman *ρ*, macro-averaged over the test antigens with at least five variants (44; 40 for retrieval), with antigen-level bootstrap confidence intervals. Every method family runs through the identical folds (Methods). As a matched reference, within-target ranking of a target’s own variants (measure-then-rank) reaches *ρ ≈* 0.37 (Section 2.1).

Across targets this ranking ability collapses (Figure 2; Extended Data Table 1). Restricted to the well-powered held-out targets (the 9 antigens with at least 30 variants, i.e. the targets for which the cross-target estimate is statistically meaningful), every method family is consistent with zero: the structure GNN reaches *ρ* = *−*0.03 [*−*0.11, +0.07], pure sequence (ESM-2) *−*0.03, gradient-boosted trees *−*0.02, *k*-nearest-neighbour retrieval +0.01, and the cofolding-confidence discriminator *−*0.01. A modern *antibody-specific* language model (AntiBERTy [11]) is no better, so the collapse is not a limitation of the sequence encoder, and predicted structure provides no cross-target advantage: the GNN sits at the wall with every other method, neither best nor worst. A macro-average over all 44 antigens with at least 5 variants is nominally higher (GNN +0.04, trees +0.15), but that is an artefact of many small, near-duplicate viral-spike constructs (*n* = 5 to 20) with unstable correlations, not genuine transfer to new targets. The cofolding-confidence model does not rank variants *within* a target either, confirming that confidence is a binder/non-binder filter, not an affinity ranker [14]. We report within-held-out-antigen Spearman; a pooled *R*^2^ is higher only because a model can predict each antigen’s mean *K_d_* offset. On named targets the collapse is explicit: HER2 (729 variants) *ρ* = *−*0.06, SARS-CoV-2 spike (262) *−*0.11, ovalbumin (88) *−*0.15, and VEGF-A (100) *−*0.05, none showing cross-target transfer.

**Figure 2:**
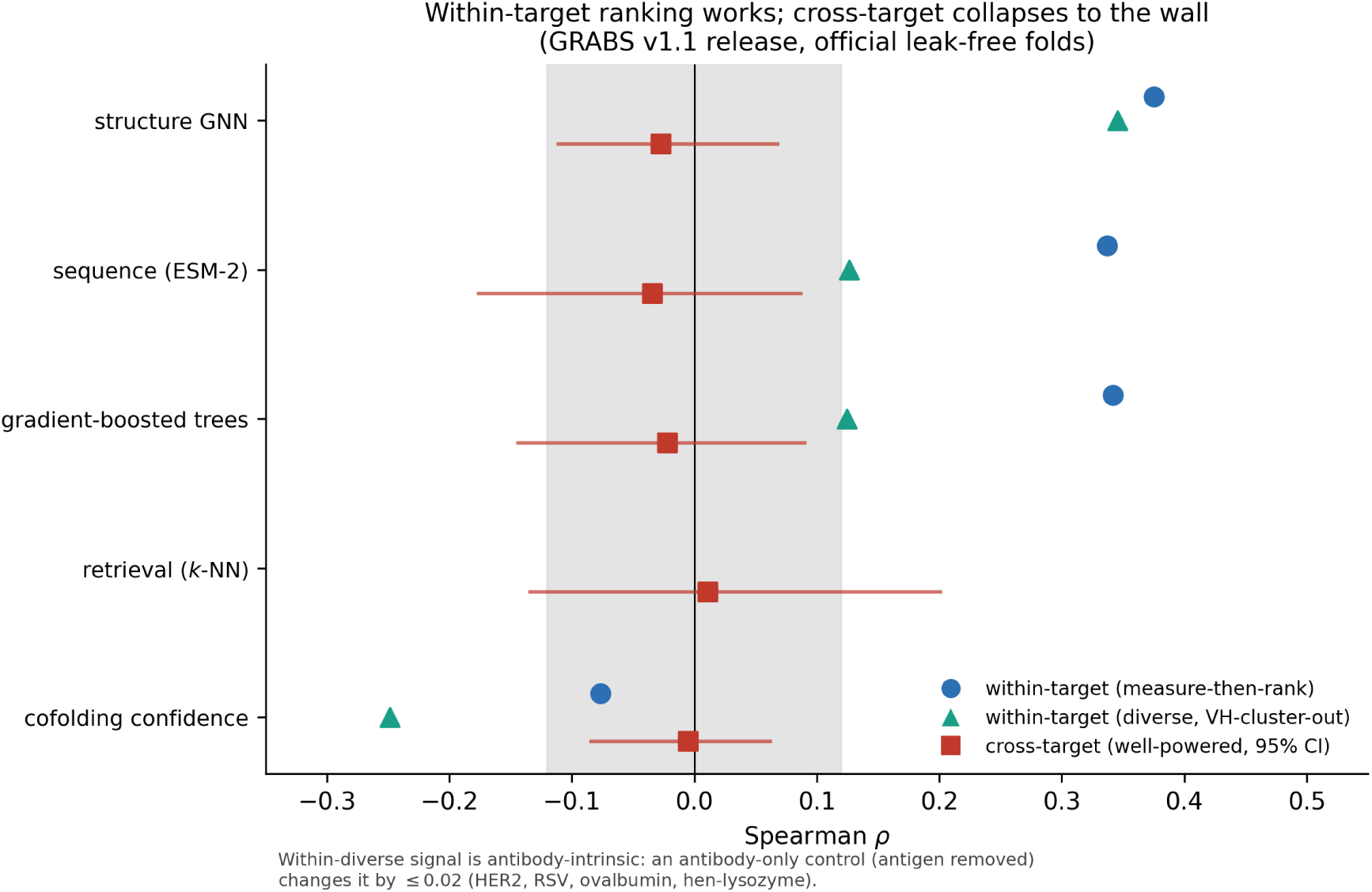
No method family generalizes affinity ranking to unseen antigens (GRABS v1.1 release, official leakage-free antigen-cluster five-fold folds). For each method: within-target measure-then-rank (blue circle; five-fold CV of a target’s own variants), within-target diverse ranking (teal triangle; leave-VH-cluster-out), and cross-target transfer (red square with 95% CI; within-held-out-antigen Spearman macro-averaged over the 9 well-powered targets). Within a target the structure GNN and feature models rank variants (measure-then-rank *ρ ≈* 0.34 to 0.38); across targets every method collapses into the wall band (*ρ ≈* 0: GNN *−*0.03, sequence *−*0.03, trees *−*0.02, retrieval +0.01). The GNN’s higher within-diverse value (0.35) is antibody-intrinsic for this model, not interface-driven: an antibody-only control (antigen nodes removed) changes it by *≤* 0.02 on HER2, RSV, ovalbumin and hen-lysozyme. The cofolding-confidence model does not rank within a target (*ρ* = *−*0.08): a binder/non-binder filter, not an affinity ranker. See Extended Data Table 1.

Measuring a target’s own variants still ranks them, but structure does not reach *within*-target affinity either. A measure-then-rank macro over sampled targets reaches *ρ ≈* 0.37, and on some diverse panels a pairwise delta-GNN ranks even VH-cluster-held-out antibodies well (hen-lysozyme 0.58, RSV 0.65). An antibody-only control settles what this signal is: dropping the antigen nodes from the graph leaves it essentially unchanged (interface contribution Δ*ρ* = *−*0.008 for HER2, +0.007 for RSV, +0.004 for ovalbumin, +0.018 for hen-lysozyme). The ranking this delta-GNN captures is therefore antibody-intrinsic, reflecting germline and composition rather than paratope-epitope complementarity (interface Δ*ρ* between *−*0.01 and +0.02). Because the same model localizes the experimental paratope only weakly (Supplementary Note 6), this bounds what this class of structural model extracts from the interface rather than proving the interface carries no signal in principle. Consistently, the largest genuinely-diverse panel (spike, 262 antibodies) collapses (*ρ* = 0.11 VH-cluster-out), while HER2’s apparent diverse signal reflects near-identical antibodies (median VH identity 0.86). Adding target-specific physics (FoldX [8]), inverse-folding likelihoods, or MM-GBSA rescoring does not change this (Extended Data Table 1). For these diverse, well-powered real targets, the within-target advantage of Section 2.1, where it exists, does not carry to any unseen antigen for any approach we tested.

The boundary is not specific to our data or models; it recurs across independent datasets and studies. On our own SKEMPI head-to-head, the structure model leads within a complex (0.548) but falls to mid-pack when an entire complex is withheld (Supplementary Note 4). On the AbRank benchmark, whose splits isolate novel antigens, a sequence-only model outperforms a hybrid sequence–structure graph network (ranking AUC 0.81 vs. 0.75) [13]. And a systematic study of equivariant ΔΔ*G* networks finds that models trained on the few hundred available experimental mutations overfit and do not generalize to complexes dissimilar from training, concluding that orders of magnitude more diverse data would be needed [14]. The same collapse appears on the most recent antibody-language-model affinity benchmark: on the six diverse antigens of AbCDR-Binding [20], within-target ranking (mean *ρ ≈* 0.66) falls to the wall across held-out antigens (*ρ ≈* 0.08), and the authors’ own released CDR-masked antibody language model—state of the art within-target—reproduces that within-target performance (*ρ ≈* 0.66) yet does not transfer to a new antigen (*ρ ≈ −*0.04; Supplementary Note 3). None of these prior studies establishes *why* the boundary exists.

### 2.3 Measuring the transfer budget: estimator and real-data validation

We now ask *why* cross-target transfer collapses. Two explanations could produce the wall: a *representational* one (predicted structures are too poor to support transfer) or an *informational* one (target-specific binding rules do not generalize even in principle). Separating them requires measuring how much of the affinity signal is transferable at all. We do so with a variance-components estimator of the cross-target transfer ceiling, *ρ*_max_ = 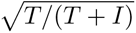 (Methods): a two-way random-effects decomposition splits the ranking-relevant affinity variance into a transferable antibody-intrinsic component *T* and a target-specific interaction *I*, and—unlike the naive sum-of-squares decomposition, which absorbs measurement noise into *I* and so understates the ceiling—it nets out a replicate-estimated noise variance and handles right-censored assays by expectation-maximization. On deterministic synthetic ground truth (Absolut! [18], where affinity is a *known, exact* function of an antigen lattice structure) the estimator recovers the noise-free ceiling to within *±*0.0003 across injected-noise levels up to twice the interaction signal, while the naive estimator degrades monotonically (Figure 3a); with replicated measurements even the naive estimator recovers, confirming the bias is a pure measurement-noise artefact.

**Figure 3:**
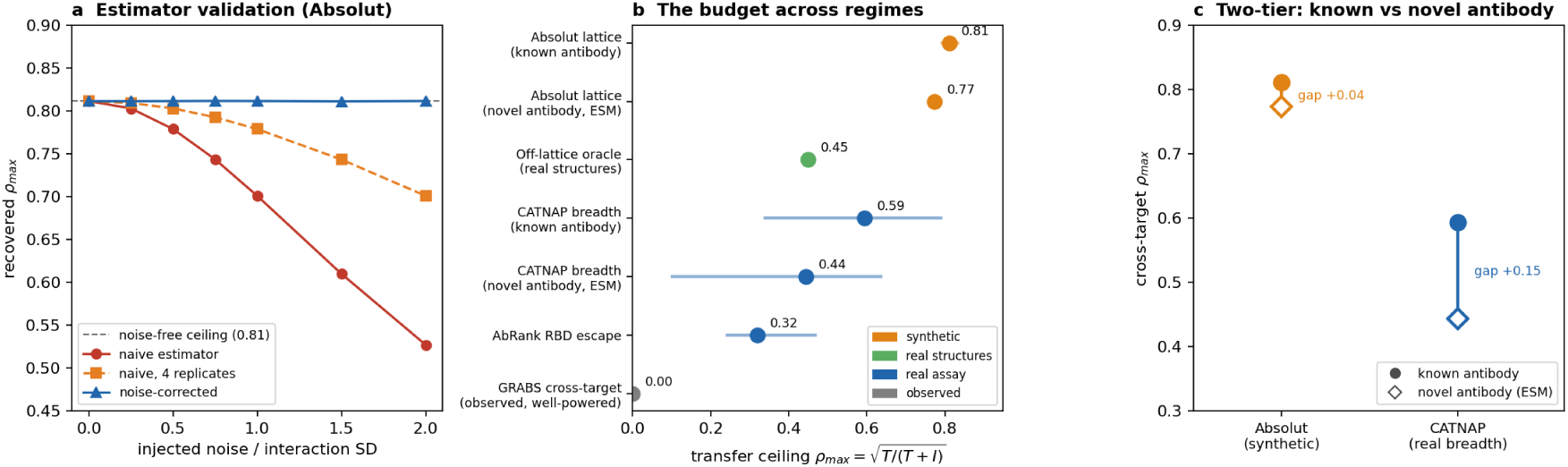
The information budget, measured across systems. **(a)** Estimator validation on the deterministic Absolut matrix: under injected measurement noise the noise-corrected estimator recovers the noise-free ceiling (*±*0.0003) while the naive sum-of-squares estimator is biased down; averaging four replicates rescues the naive estimator, confirming a pure noise artefact. **(b)** The transfer ceiling 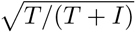 across a ladder of systems, synthetic to real (bootstrap 95% CIs where estimable). The idealized synthetic (known-antibody) ceiling is high (0.81); real structures (off-lattice oracle, 0.45) and real assays (CATNAP breadth 0.59; AbRank escape 0.32) are markedly lower; the GRABS bar is the *observed* cross-target transfer (*≈* 0 on well-powered unrelated antigens, not a ceiling, shown for reference). The synthetic oracle overstates transferability. **(c)** The two-tier structure: predicting a *novel* antibody from sequence (open diamonds, ESM-2) versus ranking a *characterized* one (filled circles). The gap is small in the deterministic synthetic system (+0.04, pure model capacity) but large on real breadth data (+0.15), quantifying the informational cost of designing a new antibody.

On a fixed, consistent panel of 86 Absolut antigens the transferable ceiling is 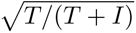 = 0.81 (95% CI 0.79–0.83), robust across antigen sub-panels. This is a *known-antibody* ceiling (the best a model can do for an antibody whose cross-target behaviour it has already observed), and it is faithful: measured directly, a main-effect predictor that transfers each antibody’s mean affinity to a held-out antigen reaches 0.806, matching the formula to within 0.008. The realistic design setting is harder, because a *novel* antibody must be predicted from sequence alone; there a strong protein language model (ESM-2) reaches 0.77 and a gradient-boosted structure model 0.72. Because Absolut’s affinity is a deterministic function of the antibody sequence, this known-versus-novel gap is model capacity, not missing information. Absolut therefore *validates the estimator and sets the idealized ceiling*, but, being deterministic, cannot by itself establish that the wall is informational rather than representational; that requires real biophysics (Figure 3c).

Real predicted structures already halve the budget. An off-lattice oracle—a dense 40*×*40 antibody– antigen matrix of 1,600 Boltz-2 complexes [5] scored by two independent structure functions (interface contacts and confidence, ipTM), with a sequence-only ESM-2 ranker blind to the structures— sits in the *same* cross-diverse-target regime as the lattice, yet its transferable ceiling is only 0.44– 0.46, about half the synthetic value (0.81). The observed cross-target ranking approaches this ceiling (0.43 [0.38, 0.48] for ipTM; 0.39 [0.35, 0.43] for contacts), rises toward it as training antigens are added (0.30 *→* 0.43 and 0.26 *→* 0.39), and collapses to zero under a permutation control that destroys the antibody*×*antigen structure (Extended Data Table 2, Extended Data Fig. 1). Because the only change from the lattice is real predicted structures in place of a synthetic one, the synthetic oracle overstates transferability—a caution for in-silico antibody engineering. (These oracle affinities are physics-based scores on predicted complexes rather than measured *K_d_*; the claim concerns the predicted-versus-observed transfer relationship, not absolute accuracy.)

The limit is confirmed, and shown to be *genuine*, on real assay panels. On CATNAP [21] (a broadly-neutralizing-antibody *×* HIV Env-strain neutralization matrix, restricted to antibodies leak-free against GRABS), the target-specific interaction is not a measurement artefact: after estimating the assay noise from replicates (*σ_є_ ≈* 0.44 in log_10_ IC_50_) and modelling the *∼* 55% right-censored (non-neutralizing) cells by expectation-maximization, the disattenuated interaction falls by only *∼* 5% and the ceiling is essentially unchanged (breadth ceiling *ρ ≈* 0.6–0.68 across block sizes). The two-tier structure reappears: transferring a *characterized* antibody to a novel strain reaches *ρ* = 0.59, but predicting a *novel* antibody from sequence alone reaches only *ρ* = 0.44 even with ESM-2—and because CATNAP is real rather than deterministic, this gap plausibly reflects a limit on how far breadth is written in sequence beyond model capacity (the two ceilings are paired on the same panel; we do not report a formal test of their difference, and their marginal intervals overlap). On AbRank RBD-mutant escape [22] (a continuous, essentially uncensored panel, so no expectation-maximization is needed) the budget is lower still, *ρ ≈* 0.32 and robust to plausible measurement noise—so the low ceiling reflects biophysical specificity, not the censoring model.

The same accounting explains the severity of the wall on GRABS itself: cross-target ranking collapses to the noise floor (well-powered Spearman *ρ ≈* 0 for every method: GNN *−*0.03, sequence *−*0.03, trees *−*0.02) against a within-target reference of *ρ ≈* 0.37, so the transferable fraction of the ranking-relevant signal is negligible, far below both the real-assay ceilings (CATNAP, AbRank) and the idealized synthetic ceiling, because real interfaces between unrelated antigens carry overwhelmingly target-specific, non-transferable information (Supplementary Note 11). The wall is therefore *informational* and, for diverse well-powered real targets, robust: it is established on real data, where the interaction survives noise and censoring correction and novel-antibody transfer is genuinely capped (HER2 *ρ* = *−*0.06, SARS-CoV-2 spike *ρ* = *−*0.11). The apparent partial transfer seen in synthetic oracles is a clean-system control, not evidence that the real wall is breakable: transfer ceilings run from 0.81 in the idealized synthetic case down to 0.32 for real escape, while the observed cross-target transfer on GRABS is *≈* 0, so even a noise-free oracle retains a large non-transferable target-specific component and overstates what real data achieve. This rules out a purely representational explanation and bounds the plausible gains from better structure prediction alone.

### 2.4 Structure helps 3D-surface properties, not sequence-dominated ones

If structure pays off through 3D geometry, it should help properties decided at a surface or interface but not those determined by the antibody sequence alone. We find a clean boundary along exactly this line (Figure 4). Evaluated as a single lumped risk, a sequence model, and the affinity GNN are all weak and statistically indistinguishable on developability (thermostability, aggregation, polyre-activity via FLAb [23] on folded Fv; Spearman between *−*0.04 and 0.13; Figure 4a), consistent with an independent benchmark that current AI models cannot reliably predict developability [24]. That view, however, lumps distinct biophysical properties and applies representations not designed for them.

**Figure 4:**
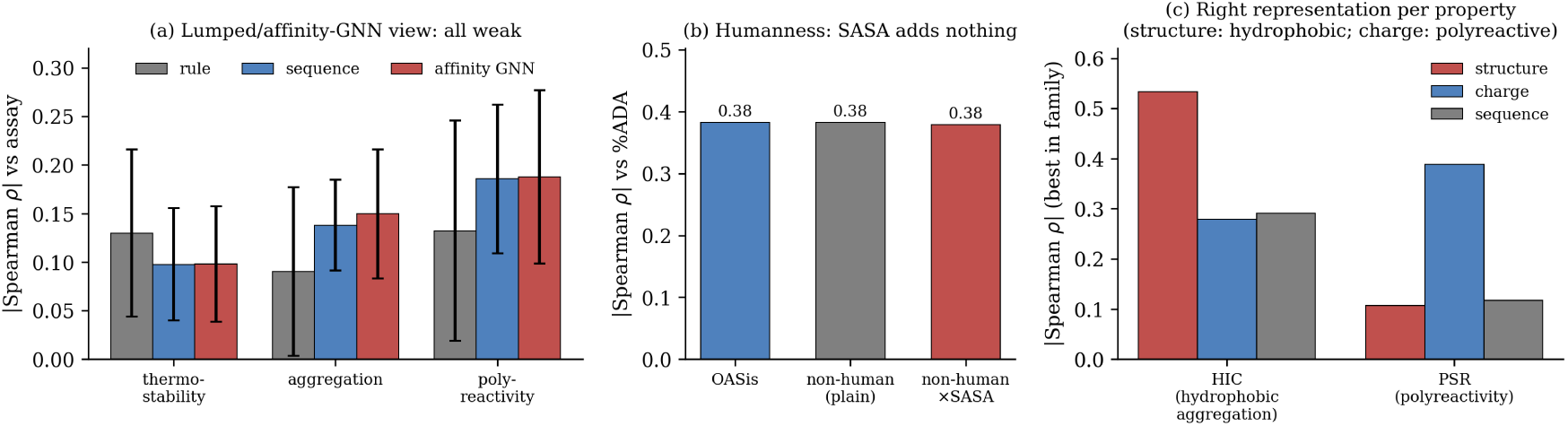
A representation-to-property boundary. (a) Treated as one lumped risk, a rule-based baseline, a sequence model, and the affinity GNN are statistically indistinguishable on thermostability, aggregation, and polyreactivity (mean *±* s.d. over five seeds), all weak. (b) Humanness is sequence-dominated: an OASis-style score and a plain non-human fraction both track clinical immunogenicity (%ADA) at *|ρ| ≈* 0.38, and weighting the non-human fraction by surface exposure (SASA) adds nothing. (c) When each property is probed with the matched representation on the Jain 2017 clinical panel, a split emerges: hydrophobicity-driven aggregation (HIC) is best predicted by a *structural* exposed-hydrophobic-surface descriptor (*ρ* = 0.53), whereas charge-driven polyre-activity (PSR) is best predicted by a *sequence* charge descriptor (*ρ* = 0.39). Structure helps this 3D-surface property through exposed-surface geometry; sequence-dominated traits do not benefit from 3D context.

When each property is instead probed with the representation matched to its physics, on the clinical-stage Jain 2017 panel [25], a sharp split appears (Figure 4c). Hydrophobicity-driven aggregation is captured by a *structural* descriptor, the solvent-exposed hydrophobic surface of the folded Fv (hydrophobic-interaction chromatography: exposed hydrophobic SASA *ρ* = 0.53, well above sequence composition 0.32 and the lumped rule *−*0.09; salting-out SGAC-SINS *ρ* = 0.45). Charge-driven self-association and polyreactivity are instead captured by a *sequence* descriptor, net or heavy-chain charge (AC-SINS *ρ* = 0.33; PSR 0.39; BVP-ELISA 0.27), with no structural gain. Structure therefore helps the 3D-surface property (hydrophobic aggregation), while charge- and sequence-encoded liabilities do not benefit from 3D context. These are not new predictors: surface-hydrophobicity and charge-symmetry descriptors underlie established developability tools [26, 27]; the contribution here is mapping representation to property.

Humanness is on the sequence-dominated side of the boundary: a sequence humanness score (OASis) [28] predicts clinical anti-drug-antibody responses (*ρ* = *−*0.38) and, across therapeutic antibodies, tracks engineering origin (*ρ* = 0.741 on Thera-SAbDab [29]), whereas weighting non-human motifs by structural solvent exposure adds nothing (Δ = *−*0.003); the conclusion is robust to the choice of metric (AbNatiV vs. OASis, *ρ* = 0.91) [30]. The unifying law is thus sharper than “structure helps only affinity”: structure helps properties set by 3D surface geometry (hydrophobic aggregation) and local structural perturbation of a known complex (single-complex mutational ranking), but it does not transfer across targets, the paratope-epitope interface itself carries no generalizable affinity signal, and sequence-dominated properties (humanness, charge-driven self-association, chemical liabilities) do not benefit from 3D context, being governed by linear sequence (for example MHC-II peptide processing) or net charge.

### 2.5 A practical consequence: measure-then-rank

The within/cross dichotomy implies a concrete deployment mode. Because affinity ranking works within but not across targets, the useful workflow is *measure-then-rank* : obtain a few experimental measurements on the target of interest, then use the model to rank the next round of candidates. In a retrospective acquisition simulation (pool of 5,000 variants, seed of 50, batches of 25 over ten rounds, ten random restarts), model-guided greedy selection recovers roughly 6*×* more of the true top-50 binders than random selection at a budget of 300 measurements (17.5 vs. 2.9; Figure 5), and the advantage grows with the measurement budget. This is the honest operating point for structure-based antibody affinity models: not zero-shot prediction for a new target, but efficient prioritization once a target has been sampled. Uncertainty-guided acquisition (upper-confidence-bound) improves further to roughly 8*×* random (23.0 vs. 2.9), its value being sample efficiency at larger budgets rather than a change in the qualitative regime; on the order of 10^2^ measured variants are needed for useful ranking, with no shortcut around this target-specific data cost.

**Figure 5:**
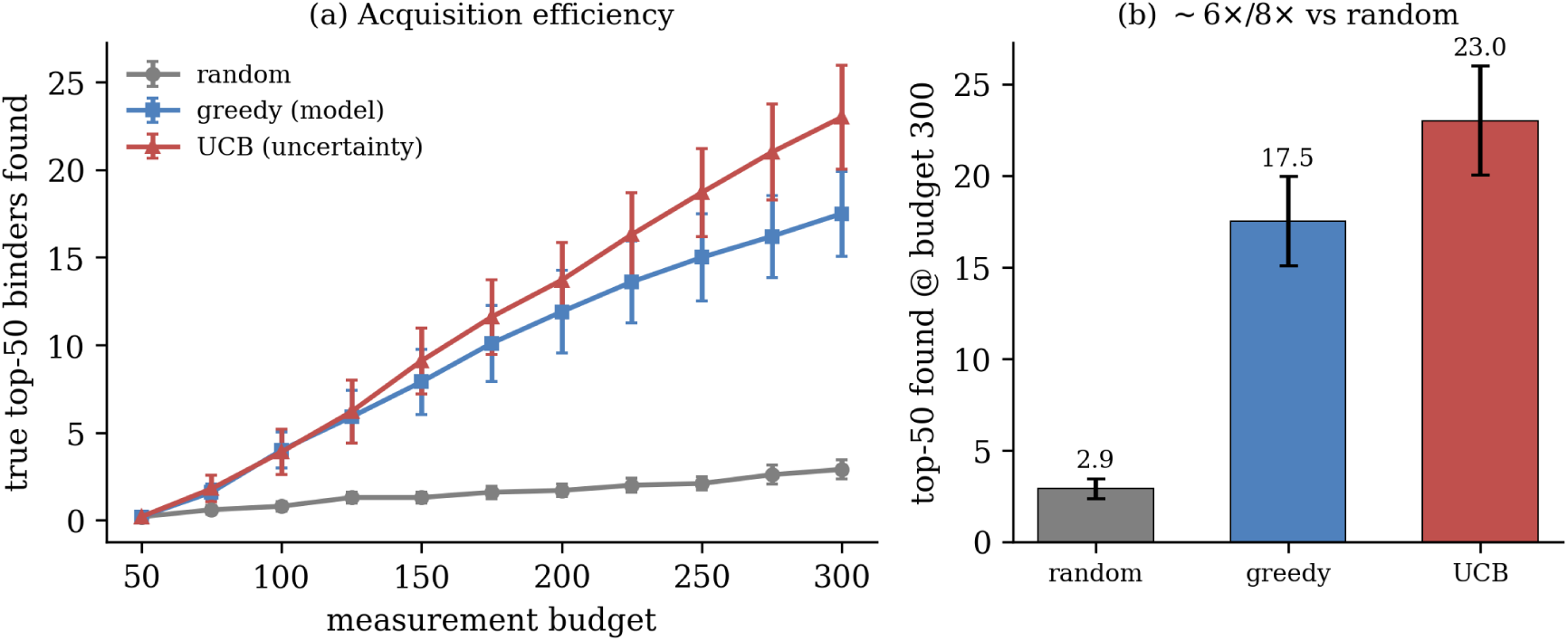
Structure-based ranking is useful within a target, given a modest number of measurements. (a) True top-50 binders recovered as a function of the measurement budget under random, greedy (model-guided), and UCB (uncertainty-guided) acquisition (mean *±* s.e.m. over ten restarts). (b) At a budget of 300, greedy and UCB recover *∼* 6*×* and *∼* 8*×* more true top binders than random selection, respectively.

The measure-then-rank claim holds on fully external data. On the AlphaSeq dataset [31] (12,000 antibodies across four lineages, measured by a yeast-display assay and 100% sequence-disjoint from GRABS), within-lineage measure-then-rank recovers the affinity ranking (mean Spearman 0.38 over the four lineages), whereas transferring a model across antibody lineages collapses (mean *≈* 0, most pairs degenerate), even though all share the same target. The within/cross dichotomy therefore reproduces on an independent, different-assay dataset the model never saw (Supplementary Note 13).

## 3 Discussion

The reach of structure in antibody engineering follows a single principle: structure helps the properties set by surface geometry and by local structural perturbation, and stops at the paratope-epitope interface. It predicts a surface-driven developability property (hydrophobic aggregation) and, within a sampled target, ranks a target’s own variants (measure-then-rank); but it fails to generalize to new targets, and a controlled ablation shows that even within a target the antibody-antigen interface adds nothing over the antibody alone for this structural model, so the within-target signal it captures is antibody-intrinsic. It does not help sequence-dominated properties (humanness, charge-driven self-association). We identify the cause by *measuring* the transfer budget with a noise-corrected estimator across a ladder of synthetic and real systems: the cross-target ceiling is set by a target-specific interaction that, on real neutralization and escape panels, survives noise and censoring correction, so it is an informational limit rather than an artefact of imperfect predicted structures. It is attenuated but not removed by target diversity, and synthetic oracles overstate how far it can be crossed.

### Relation to concurrent work

That cross-target affinity models degrade under leakage-free splits is now reported independently: Hummer et al. find equivariant ΔΔ*G* networks are not robust to train–test cutoffs [14], the AbRank benchmark documents poor transfer to held-out antigens [13], a universal-benchmark evaluation finds sequence models match or exceed structure-based predictors on held-out antigens [32], an SE(3)-equivariant model generalizes only modestly under stringent antigen-sequence splits [33], and the effect echoes the memorization long known for protein–ligand affinity [12]; and even large domain-specific and foundation models—AlphaBind, pre-trained on 7.5 million quantitative measurements [34], and the multi-modal MAMMAL [35]— still require per-target fine-tuning and do not generalize zero-shot across binding landscapes, the very ceiling our information budget quantifies. Our contribution is orthogonal. That body of work attributes the collapse to *data scarcity* —an extrinsic limit expected to yield to more and more-diverse data [14]—whereas we measure a residual ceiling (quantified as an information budget with a noise-corrected, censoring-aware estimator, and real on the neutralization and escape panels) that survives with the antigen structure known *exactly* and does not close as training antigens are added without limit. The two views are not in conflict: our analysis targets precisely the cross-antigen transfer that remains *after* data scarcity is removed, and shows this residual to be informational rather than representational, so more or better-predicted structures alone cannot close it even as more diverse targets attenuate it. Our use of the Absolut! oracle [18] is likewise complementary to Ursu et al. [36], who use it to study how negative-set composition shapes learned binding rules; we instead exploit known-exact structure to separate representational from informational causes of the cross-target ceiling.

Four implications follow. First, it bounds the expected return on better structure prediction alone for cross-target affinity, which may be necessary but is not sufficient. Second, it makes the case that progress on the cross-target problem will come from *more diverse targets* (foundation models trained across many antigens) and from *real physics* (free-energy methods that model the target-specific interaction directly), rather than from architectural changes. Third, it argues that the community should retire leakage-prone cross-target metrics in favour of antigen-grouped evaluation, and should report the within/cross distinction explicitly. Fourth, because synthetic oracles overstate transferability, cross-target claims should be calibrated against real crossed panels, not synthetic ground truth alone.

### The generative-design turn

The field’s fastest-moving response is not better cross-target *prediction* but de novo *design*: generative binder and antibody models (AlphaProteo [37]; RFdiffusion-designed antibodies [38]) and closed-loop optimization (lab-in-the-loop [39]) now yield epitope-specific binders with experimentally validated nanomolar affinities. This is consistent with, not contrary to, our result: design-then-test *sidesteps* the cross-target prediction wall—it never asks a model to rank an unseen antibody against an unseen target—which is exactly the measure-then-rank operating point we advocate, where progress comes from spending experimental measurements rather than from zero-shot transfer. The information budget clarifies two features of this era. First, generative hit rates remain strongly epitope-dependent and hard to predict a priori (from near zero to a majority of designs, across methods), because the designable signal is bounded by the same target-specific information. Second, ever-more-accurate structure and interface engines (e.g. AlphaFold-3-scale antibody modelling [40]) improve within-target ranking and design fidelity but cannot, by our analysis, cross an *informational* cross-target boundary—better structures are necessary but not sufficient. The budget thus reframes the design era, predicting where transfer will fail and how much measurement each target demands.

The reach principle collapses into a simple decision rule for practitioners (Figure 6): use structure only when the property is decided at a 3D surface or interface *and* the target has been sampled; otherwise rely on sequence or charge descriptors, and obtain affinity by measure-then-rank. Every property we examined fits this rule.

**Figure 6:**
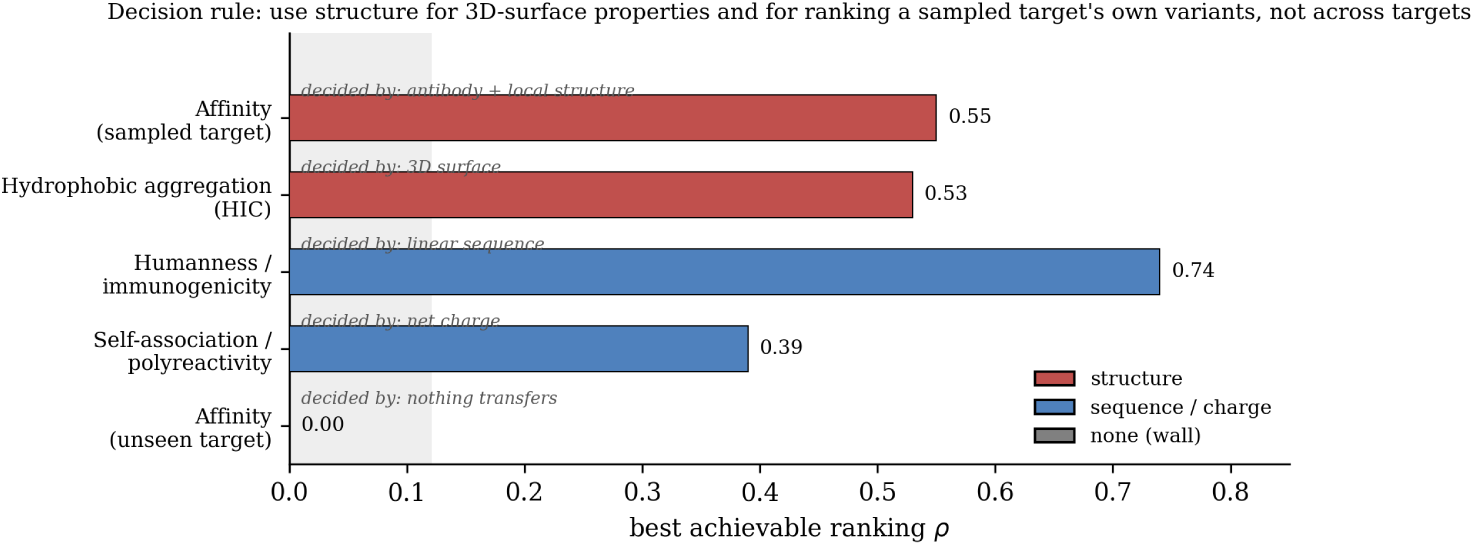
A decision rule derived from the reach principle. For each antibody-engineering task, the bar shows the best achievable ranking Spearman *ρ* and its colour the winning representation: structure (red; affinity for a sampled target and hydrophobic aggregation), sequence/charge (blue; humanness and charge-driven self-association/polyreactivity), or none (grey; affinity for an unseen target, the informational wall). Structure is the right representation exactly when the property is set by 3D surface or interface geometry *and* the target is sampled; otherwise sequence/charge descriptors and measure-then-rank are the honest choice. The humanness bar shows the best achievable value, *ρ* = 0.74 (engineering origin on Thera-SAbDab); clinical immunogenicity (%ADA) is weaker at *ρ* = 0.38. All rows are instantiated by results in this paper.

The study is retrospective: it characterizes existing data and models rather than validating predictions prospectively in the laboratory, and its real-data evidence comes from retrospective crossed panels (neutralization, escape) rather than a designed prospective test. The synthetic lattice serves only to validate the estimator and set an idealized ceiling—its magnitudes do not map directly to real binding, which is why the informational claim rests on the real panels. Transfer ceilings are regime- and panel-dependent, and we therefore report them as ranges with confidence intervals rather than single values. Finally, the public landscape imposes a label-diversity ceiling: no current corpus is simultaneously large, clean, and broadly target-diverse in quantitative *K_d_*, and no real crossed panel spans genuinely unrelated targets—itself part of why the wall is hard to cross with data alone.

## 4 Methods

### Data

Affinity data are drawn from GRABS, a curated, content-addressed corpus of 4,380 antibody–antigen records with experimental *K_d_* across 2,015 antigens, with predicted complex structures, per-residue ESM-2 embeddings, and geometric graphs, released as a companion data descriptor (GRABS v1.1) [15]; 4,282 records are structure-backed and 4,020 form the leak-free benchmark used here, evaluated under the release’s official antigen-cluster five-fold split (549 clusters). Mutational ΔΔ*G* data are from SKEMPI 2.0 [16] and AB-Bind [17]; crystal-structure benchmarking uses AbBiBench [19]; and synthetic ground truth is from Absolut! [18]. Antigen sequences are clustered with MMseqs2 [41] for leakage-free grouping; structures are predicted with Boltz-2 [5]; residue features use ESM-2 [10]; numbering uses ANARCI [42].

### Affinity model

The structure model is a delta-siamese geometric-vector-perceptron (GVP) graph neural network [6]. Each residue node carries its 1,280-dimensional ESM-2 embedding concatenated with a one-hot IMGT region label, chain identity, and solvent accessibility; edges encode sequential adjacency and spatial contacts. GVP layers propagate messages SE(3)-equivariantly; antibody and antigen subgraphs are combined by a bilinear cross-attention block whose weights provide the interpretability maps (Supplementary Note 6); node states are pooled per CDR and framework region; and a multilayer-perceptron head predicts log_10_ *K_d_*. For mutational tasks the network is applied in a siamese configuration that predicts the difference between variant and wild type (ΔΔ*G*). Training minimizes mean-squared error with the Adam optimizer and early stopping under the antigen-grouped folds; the full architecture and hyperparameters are in the released code.

### Baselines

FoldX [8] computes ΔΔ*G* on both crystal and Boltz-2 predicted complexes; GearBind is evaluated with and without contrastive pretraining [7]; mCSM-AB2 [9] is included with an applicability and leakage analysis (Supplementary Note 2); a sequence baseline mean-pools ESM-2 embeddings into a matched-capacity multilayer perceptron; gradient-boosted trees use aggregated sequence, physicochemical, and structure-confidence features; cofolding combines Boltz-2 with a learned binder discriminator; inverse-folding likelihoods and nearest-neighbour retrieval provide training-free, structure-aware baselines; and MM-GBSA rescoring is computed with OpenMM.

### Evaluation

Within-target performance is measured within complexes. Cross-target performance uses leave-one-antigen-out over MMseqs2 antigen clusters (min. sequence identity 0.8, coverage 0.8; 1,752 antigens *→* 549 clusters), so that near-duplicate antigens are held out together and cannot leak across folds; the reported metric is the within-held-out-antigen Spearman *ρ* macro-averaged over test antigens with *≥* 5 variants. We report Spearman *ρ* and *R*^2^ with 95% bootstrap confidence intervals (*B* = 1,000); model differences are assessed with paired-*t*/Wilcoxon tests and Benjamini–Hochberg correction across comparisons.

### Absolut! ground-truth experiments

Absolut! antigens are parsed from the lattice library to recover the exact structure; an interaction-capable model receives the true-structure descriptor plus the CDR-region features, with a leakage-free CDR split and a fixed held-out antigen set. The structure contribution is isolated by ablating the antigen descriptor, and cross-target performance is measured as a function of the number of training antigens.

### Transfer-ceiling estimator

The cross-target ceiling *ρ*_max_ = 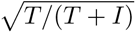 is estimated from a crossed antibody*×*antigen value matrix by a two-way random-effects variance decomposition: *T* (antibody main-effect variance) and *I* (antibody*×*antigen interaction) are recovered from the expected mean squares, the measurement variance *σ*_ε_^2^ is estimated from within-cell replicates and subtracted from *I* (disattenuation), and right-censored assays (e.g. CATNAP IC_50_ above the assay limit) are handled by an expectation-maximization scheme propagating the first and second truncated-normal moments. The estimator is validated on the deterministic Absolut! matrix by injecting Gaussian noise of known magnitude (as a multiple of the interaction SD) and confirming recovery of the noise-free ceiling; confidence intervals are by bootstrap over antibodies and antigens, and ceilings are reported across antigen panels and block sizes to expose panel dependence. The two-tier (known vs. novel antibody) ceiling contrasts a main-effect oracle—each antibody’s mean transferred to a held-out target—with a sequence-only ESM-2 [10] model trained on a disjoint antibody split (leave-one-antigen-out over targets). Real crossed panels are CATNAP [21] (a broadly-neutralizing-antibody *×* HIV Env-strain neutralization matrix) and the RBD-mutant escape subset of AbRank [13], from the deep mutational scan of [22], each restricted to antibodies with *<* 90% VH identity to any GRABS antibody (leakage-free); potency is *−* log_10_ IC_50_ and escape is the log binding-retention ratio.

### Intrinsic properties

Developability (thermostability, aggregation, polyreactivity) uses FLAb [23] with Boltz-folded Fv; humanness uses OASis [28], validated against clinical anti-drug-antibody responses and engineering origin, and cross-checked with AbNatiV [30]. The representation-to-property analysis (Supplementary Note 10) uses the clinical-stage Jain 2017 panel [25] with Fv structures from ABodyBuilder2 [43] and solvent-accessible surface area from a Shrake–Rupley estimator; structural (exposed-hydrophobic surface) and sequence (net/heavy-chain charge) descriptors are compared per biophysical readout by single-descriptor correlation and five-fold cross-validation. The measure-then-rank simulation performs greedy and uncertainty-guided acquisition on held-out target variants.

## Data availability

The GRABS dataset and derived artifacts are available at [15]; benchmark datasets are from their original sources [16, 17, 18, 23].

## Code availability

All models, baselines, evaluation, and the transfer-budget analysis are released as deterministic, parameterized scripts with the dataset repository [15].

## Author Contributions

L.M. designed the study, implemented the models and experiments, and wrote the manuscript. J.M. and P.P.Z. supervised the work and revised the manuscript.

## Competing Interests

The authors declare no competing interests.

## Extended Data

**Extended Data Table 1:** Within-vs cross-target ranking for five method families on the GRABS v1.1 release (4,020 benchmark records, 1,752 antigens, 549 clusters), under the release’s official leakage-free antigen-cluster five-fold split. Within-target is shown in two regimes: measure-then-rank (MTR; a target’s own variants, random five-fold) and diverse (leave-VH-cluster-out). Cross-target is the within-held-out-antigen Spearman macro-averaged over well-powered targets (*n ≥* 30 variants; 9 antigens); the all-antigen macro (*n ≥* 5; 44 antigens) is shown for comparison and is inflated by small near-duplicate constructs. Across targets every family is consistent with zero. The structure GNN’s higher within-diverse value (0.35) is antibody-intrinsic: an antibody-only control (antigen nodes removed) changes it by *≤* 0.02 on HER2, RSV, ovalbumin, and hen-lysozyme, so for this model the interface adds nothing over the antibody alone. The cofolding-confidence model does not rank within a target (*ρ* = *−*0.08): a binder/non-binder filter, not an affinity ranker. Physics and likelihood baselines (FoldX, inverse-folding, MM-GBSA) are evaluated in their applicable regimes in Supplementary Note 1. Reproducible from the release via `scripts/recompute_fig2_v22.py--release`.

| **Method family** | **Within (MTR)** | **Within (diverse)** | **Cross** (*n* ≥ 30) **[95% CI]** | **Cross (all)** |
|---|---|---|---|---|
| Structure GNN (delta-siamese) | +0.38 | +0.35 | −0.03 [−0.11, +0.07] | +0.04 |
| Sequence / ESM-2 | +0.34 | +0.13 | −0.03 [−0.18, +0.09] | +0.06 |
| Gradient-boosted trees | +0.34 | +0.13 | −0.02 [−0.14, +0.09] | +0.15 |
| Retrieval (*k*-NN) | — | — | +0.01 [−0.13, +0.20] | −0.02 |
| Cofolding discriminator | −0.08 | −0.25 | −0.01 [−0.08, +0.06] | −0.07 |

**Extended Data Fig. 1:**
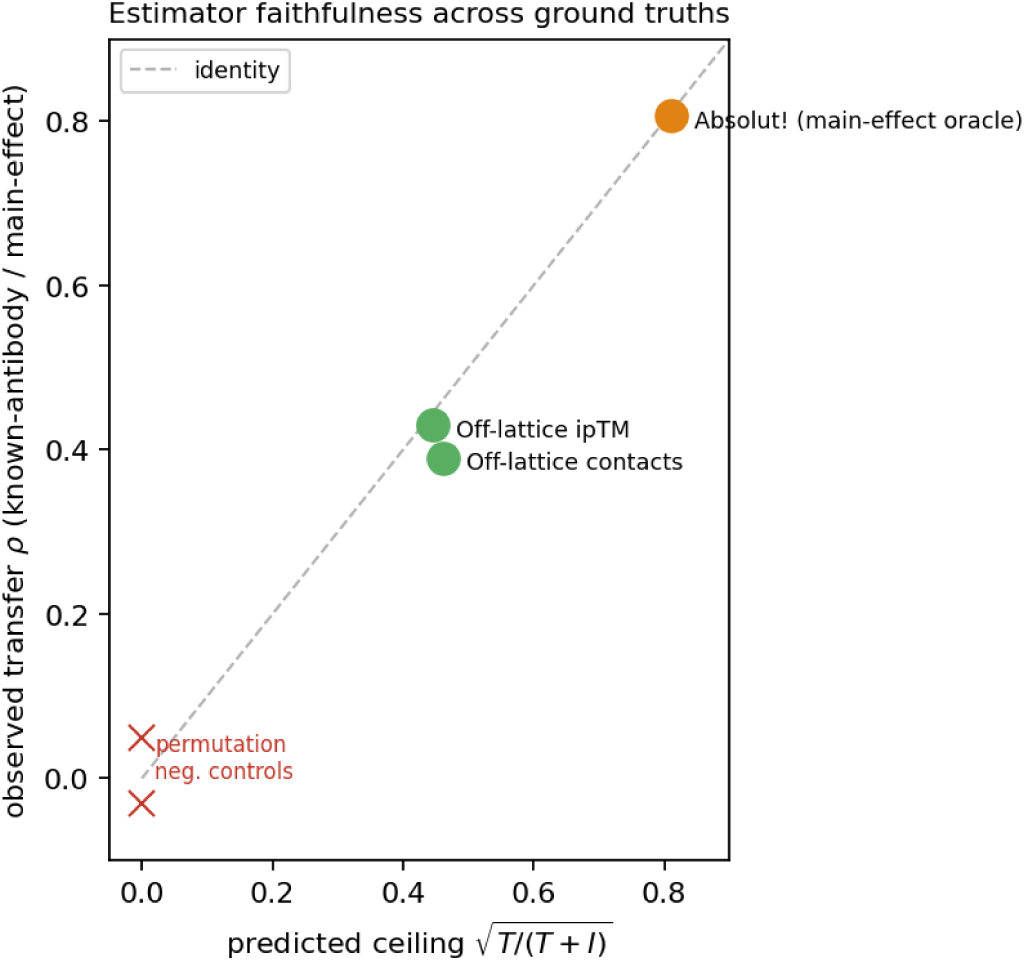
The transfer-ceiling estimator is faithful across independent ground truths. Predicted ceiling 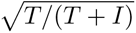 (x) versus the empirically-observed transfer of a known-antibody / main-effect ranker (y): for Absolut! the formula (0.81) matches a direct main-effect oracle (0.806); for the off-lattice oracle the predicted ceilings (0.44–0.46) match a sequence-only leave-one-antigen-out ranker (0.39–0.43). Points lie near the identity line; permutation negative controls collapse to the origin. The estimator therefore recovers the achievable transfer across ceilings spanning 0.44 to 0.81; the residual is the non-transferable, target-specific interaction, genuine on real assays (Fig. 3).

**Extended Data Table 2:**
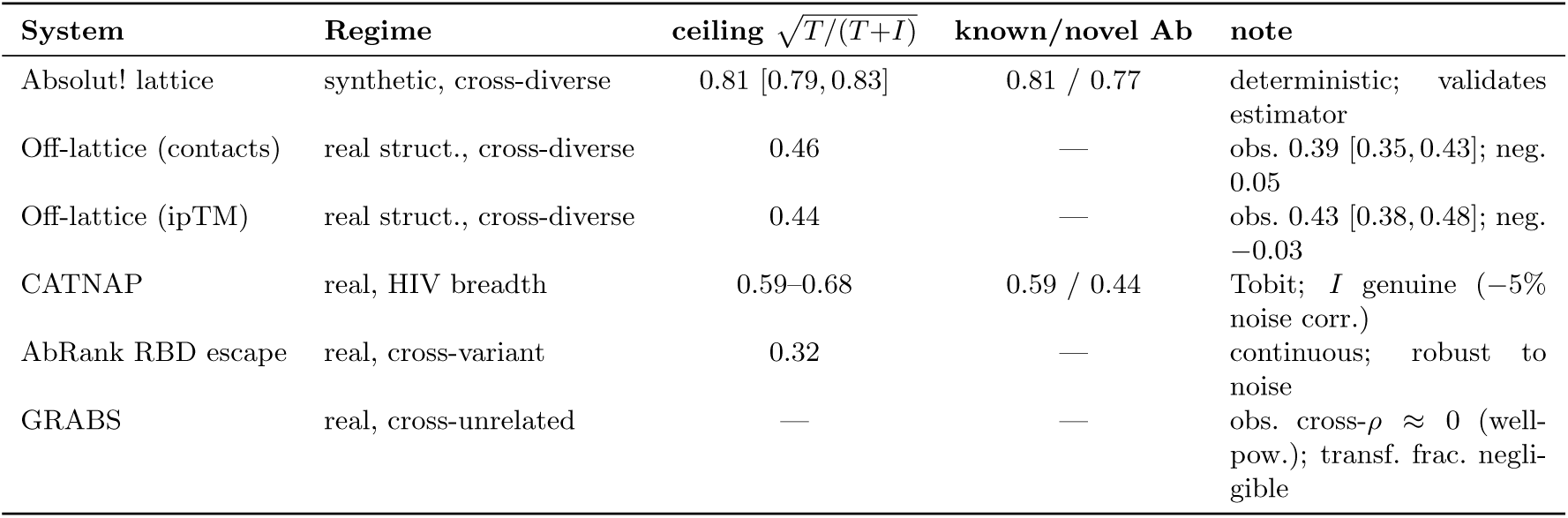
The information budget across systems, from the noise-corrected, censoring-aware variance-components estimator (Methods). The transferable ceiling 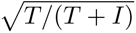 falls from an idealized synthetic value (0.81; deterministic Absolut!, which validates the estimator) through real predicted structures (off-lattice oracle, *≈* 0.45) to real assays (CATNAP breadth 0.59– 0.68; AbRank escape 0.32); on real panels the interaction is genuine, surviving noise and censoring correction. Where a sequence-only ESM-2 leave-one-antigen-out ranker is available, the observed cross-target *ρ* approaches the predicted ceiling and collapses under a permutation negative control (off-lattice: scaling over 5–39 antigens). “Known/novel Ab” is the two-tier ceiling: ranking a characterized antibody vs. predicting a novel one from sequence (the design setting). Ceilings are panel-dependent and reported as ranges/CIs.

| **System** | **Regime** | **ceiling** $\sqrt{T/(T+I)}$ | **known/novel Ab** | **note** |
|---|---|---|---|---|
| Absolut! lattice | synthetic, cross-diverse | 0.81 [0.79, 0.83] | 0.81 / 0.77 | deterministic; validates estimator |
| Off-lattice (contacts) | real struct., cross-diverse | 0.46 | — | obs. 0.39 [0.35, 0.43]; neg. 0.05 |
| Off-lattice (ipTM) | real struct., cross-diverse | 0.44 | — | obs. 0.43 [0.38, 0.48]; neg. −0.03 |
| CATNAP | real, HIV breadth | 0.59–0.68 | 0.59 / 0.44 | Tobit; *I* genuine (−5% noise corr.) |
| AbRank RBD escape | real, cross-variant | 0.32 | — | continuous; robust to noise |
| GRABS | real, cross-unrelated | — | — | obs. cross-*ρ* ≈ 0 (well-pow.); transf. frac. negli-gible |

## Supplementary Information

### Supplementary Note 1

#### Physics and likelihood baselines

FoldX, inverse-folding likelihood, and MM-GBSA are not affinity rankers trained on GRABS; because they require per-complex structures and per-mutation energetics, we evaluate each in its applicable regime rather than at GRABS scale. FoldX ΔΔ*G* ranks a data-rich affinity-maturation lineage at *ρ* = *−*0.001 (*n* = 108) and, across distinct targets, gives *ρ* between *−*0.23 and 0 (near zero); it further collapses on predicted rather than crystallographic structures (Section 2.1). Inverse-folding likelihood (ProteinMPNN) scored zero-shot on complex backbones yields cross-target *ρ* = *−*0.03 on average (range [*−*0.29, 0.15] over eight held-out targets). MM-GBSA rescoring reaches *ρ ≈* 0.56 *within* a single data-rich lineage but is a within-target tool requiring the target’s own structures. None of the three recovers a cross-target signal, consistent with the five uniform families above and with independent benchmarks [13, 14].

### Supplementary Note 2

#### mCSM-AB2 has no fair arena

mCSM-AB2 [9] scores only single-point substitutions, but the affinity benchmarks are *∼*99% multi-point mutants, so it is inapplicable to most entries; moreover, all antibody complexes in the head-to-head appear in its training set, so its apparent accuracy (*ρ ≈* 0.94) reflects memorization rather than generalization.

### Supplementary Note 3

#### Cross-target wall on the AbCDR-Binding benchmark (Talaei et al.)

We re-analyzed the six antigen panels of AbCDR-Binding [20] under a leakage-free protocol (fresh five-fold cross-validation within each panel for the within-target estimate; leave-one-antigen-out across the six panels, with per-panel *z*-scored labels, for cross-target). Our sequence families reach the same within-target regime reported by the authors (ESM-2 ridge mean *ρ* = 0.66; range 0.55–0.82; a random within-panel split that, as noted above, inflates single-mutant estimates), which collapses across held-out antigens (mean *ρ* = 0.08). Running the authors’ *own* released model (AbCDR-ESM2) with their exact embedding-plus-ridge recipe reproduces their within-target numbers (e.g. anti-fluorescein 0.72, anti-H1 0.75, matching their reported 0.73/0.76) and likewise collapses across antigens (mean *ρ* = *−*0.04), confirming the boundary on a competitor’s benchmark and model. This is a transfer test, not a within-target method comparison. A methodological caveat we note in passing (and treat fully elsewhere): on single-mutant panels, random *k*-fold splits leave one-mutation-apart near-duplicates across folds and inflate the within-target estimate—a position-held-out split lowers it substantially (e.g. anti-lysozyme 0.58 *→* 0.21).

### Supplementary Note 4

#### SKEMPI two-scenario head-to-head

Within-complex (measure-then-rank): our model 0.548 *>* GearBind+pretrain 0.465 *>* FoldX 0.436 *>* GearBind 0.404. Cross-complex (entire complex withheld): the within-complex ranking of our model falls to 0.42, mid-pack among the off-the-shelf baselines (FoldX 0.44, GearBind 0.40, GearBind+pretrain 0.47); on the pooled metric across all variants it drops to 0.35, below GearBind (0.47–0.54). The win is confined to the within-target regime.

### Supplementary Note 5

#### Structure quality controls the physics baseline

On AbBiBench crystal complexes FoldX attains mean *|ρ|* = 0.11 vs. the structure GNN’s 0.42 on the same crystal inputs. On predicted (Boltz-2) structures FoldX collapses further to *ρ ≈* 0 (data-rich lineage, *n* = 108), isolating structure quality as the driver.

### Supplementary Note 6

#### Interpretability

Cross-attention weights recover the experimental paratope only weakly (AUC *≈* 0.56, below a CDR-membership baseline), and per-residue saliency does not reliably localize affinity-driving residues either (AUC *≈* 0.43, near chance). We therefore make no strong interpretability claim: the model’s attributions do not cleanly recover a static paratope.

### Supplementary Note 7

#### Predicted vs crystal structures

On the 32 complexes for which both a Boltz-2 prediction and a crystal structure are available, predicted and crystal interfaces agree for roughly three-quarters, while about a quarter differ substantially — a caveat for any structure-dependent result computed on predicted rather than crystallographic complexes.

### Supplementary Note 8

#### Parameter-efficient fine-tuning does not lower the data cost

LoRA/PEFT adaptation does not reduce the number of target measurements required for useful ranking, confirming there is no transfer-learning shortcut around the target-specific label budget.

### Supplementary Note 9

#### Absolut! specificity diagnostic and panel dependence

Over 6.9M shared antibodies the mean pairwise cross-antigen Spearman is 0.41. The two-way decomposition is *panel-dependent* : an early 8-antigen subset gave a ceiling of 0.71, but on a fixed, consistent 86-antigen panel (and robustly across sub-panels, 0.79–0.83) the ceiling is 0.81; we therefore report the 86-antigen value and treat the ceiling as a panel-dependent range rather than a single number. Estimator choice matters less than panel here (sum-of-squares and variance-components agree to within 0.01 on this deterministic matrix), because Absolut! is noise-free; the noise correction becomes load-bearing only on real assays. Transferable and target-specific components are estimated empirically, not imposed.

### Supplementary Note 10

#### Representation-to-property mapping for developability

On the clinical-stage Jain 2017 panel (*n* = 137 antibodies), Fv structures were predicted with ABody-Builder2 [43] and solvent-accessible surface area computed with a Shrake–Rupley estimator; we correlated single physically-interpretable descriptors (no training) and five-fold cross-validated linear models with each biophysical readout. Hydrophobicity-driven readouts are predicted by *structural* exposed-hydrophobic surface: HIC retention, exposed hydrophobic SASA *ρ* = 0.53 (five-fold CV 0.58 *±* 0.02) vs. sequence composition 0.32 and the lumped liability rule *−*0.09; SGAC-SINS *ρ* = 0.45. Charge-driven readouts are predicted by *sequence* charge and show no structural gain: AC-SINS (self-association) net charge *ρ* = 0.33; PSR (polyreactivity) heavy-chain charge 0.39; BVP-ELISA net charge 0.27. These descriptors are standard biophysical quantities underlying existing developability tools [26, 27]; the point is the mapping of representation to property, not a new score. Structures are ABodyBuilder2 predictions rather than crystals, which makes the analysis deployable from sequence alone.

### Supplementary Note 11

#### The cross-target wall as an information budget

For a target-blind model the achievable cross-target ranking is bounded by the transferable fraction of the ranking-relevant variance, *ρ*_max_ = 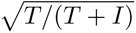, with *T* (transferable antibody-intrinsic) and *I* (target-specific interaction) from a two-way random-effects decomposition; unlike a sum-of-squares split, the estimator nets out a replicate-estimated measurement variance and handles right-censored assays by expectation-maximization (Methods). *Validation:* on the deterministic Absolut! Matrix it recovers the noise-free ceiling to *±*0.0003 under injected noise up to twice the interaction signal, whereas the naive estimator is biased down (e.g. 0.81 → 0.70 at noise equal to the interaction SD). Synthetic: on a fixed 86-antigen Absolut panel 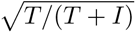 = 0:81 (CI 0:79-0:83), matching a directly-measured known-antibody transfer oracle (0.806); a novel antibody predicted from sequence reaches 0.72–0.77 (the gap is model capacity, as Absolut is deterministic). *Real:* the budget is smaller and not a noise artefact: CATNAP breadth 0.59–0.68 with the interaction surviving noise (*σ_є_ ≈* 0.44) and censoring (*∼* 55%) correction (interaction *−*5%), two-tier 0.59 (known) vs. 0.44 (novel); AbRank escape 0.32; and on GRABS cross-target ranking is *≈* 0 (well-powered Spearman), a negligible transferable fraction. The wall is thus a property of the information available (regime-dependent, and overstated by synthetic oracles), not of the model class.

### Supplementary Note 12

#### Repertoire-invariance of the synthetic transfer plateau

The synthetic result does not depend on the antibody repertoire used to build it. Repeating the experiment with a *human* CDR3 repertoire (Absolut! human bindings, 18 antigens) reproduces the pattern: within-target *ρ ≈* 0.90 versus a cross-target plateau of *ρ ≈* 0.76, close to the murine plateau (*≈* 0.72). This plateauing transfer is a property of the synthetic antigen–antibody oracle, not of the murine repertoire in which we first measured it; these are known-antibody transfer plateaus on the deterministic oracle, a clean-system control, not the informational ceiling itself (Section 2.3). They overstate what transfers on real data (Section 2.2), where diverse well-powered targets show no transfer (HER2 *ρ* = *−*0.06, spike *ρ* = *−*0.11).

### Supplementary Note 13

#### External validation on AlphaSeq

As an out-of-distribution check, we used the MITLL/AAlphaBio AlphaSeq dataset [31]: 12,000 antibodies across four lineages (AAYL49–52) against a common target, measured by a yeast-display assay unrelated to the SPR/ELISA *K_d_* in GRABS, and 100% *sequence-disjoint* from GRABS (0/48,928 heavy chains shared). This is the closest computational proxy to a prospective test: the corpus was never seen and the assay differs. Within-lineage measure-then-rank (five-fold CV on AntiBERTy embeddings) recovers the affinity ranking with mean Spearman 0.38 (per lineage 0.24–0.46), confirming that the deployment mode generalizes to independent, out-of-assay data. Transfer across lineages collapses (mean *≈* 0; 8*/*12 pairs degenerate to near-constant predictions), reproducing the within/cross dichotomy: ranking is specific to the sampled family, not transferable.

## Notes

### Competing Interest Statement

The authors have declared no competing interest.

### Funding

- Sao Paulo Research Foundation: 2024/01200-8

